# A Network Polypharmacological Approach to Combinatorial Drug Repurposing for Diffuse Intrinsic Pontine Glioma

**DOI:** 10.1101/2020.06.14.150714

**Authors:** Finlay MacLean, Javad Nazarian, Justyna Przystal, Pan Pantziarka, Jabe Wilson

**Affiliations:** University Children’s Hospital Zurich

## Abstract

Despite five decades of clinical investigations, there is currently no effective treatment for children diagnosed with Diffuse Intrinsic Pontine Glioma (DIPG). We now understand that DIPGs share the same histone 3 mutation and fatal prognosis as other diffuse midline gliomas (DMGs), which led to the introduction of a new entity referred to as DMG, H3 K27M mutant. Indeed, therapeutics indicated for other brain neoplasms have proven ineffective for DIPGs. We posit that by using a polypharmacological approach to determine drug combinations that target distinct mechanistic pathways of DIPG, it is more likely that an efficacious treatment will be developed. We predict monodrug therapies using a link prediction model trained on various embeddings of a drug-disease regulatory network and physicochemical properties of small molecules and proteins. We validate the *in silico* predictions by performing cell viability assays on patient-derived cell cultures for notable therapeutics. Using FDA-approved drugs as a proxy for viability of a drug pair for combinatorial use, we develop a model to predict the synergism of the relationship between drug pairs. Finally, we calculate the transitive probability that a drug pair contains drugs that individually regulate DIPG, are blood-brain barrier penetrant, and the drug pair are suitable for combined use. We find only moderate agreement between computational predictions and experimental results for both monodrug and multidrug therapies, we believe due to the heterogeneity of the disease, the difficulties of modelling brain permeability, and an inherent literature bias in the knowledge graph. Such challenges need to be overcome to develop an efficacious therapy for this disease.

## I. Introduction

To date, there is no approved and satisfactory means of treatment for DIPG. Repurposed drugs hold great promise as potential treatments; namely due to their established safety profiles and manufacturing routes, minimising time to use these drugs at the point-of-care [1]. Whilst there have been over 200 clinical trials for DIPG [2], none have shown great clinical promise. This is in part due to a lack of knowledge of the pathophysiology of the disease, lack of effective drugs targeting tumorigenic pathways, and difficulties in drug delivery due to the blood-brain barrier.

Systems pharmacology and network medicine (NM) approaches to drug discovery and drug repurposing have proved efficient methods to highlight potential drug candidates [3], [4]. NM treats biological networks as heterogeneous information systems; correlating network topology and node properties with biological processes, functions, pathways and interactions. From a systems biology point of view, a disease can be seen as a selection of genes within a network, whose misregulation culminates in changes in biological processes and pathways. Similarly, drugs can be modelled by their drug targets, and the propagatory effect that their perturbance has upon the network. The aim of NM is to identify drugs and diseases in which the network perturbance of the disease state is reversed by the perturbance of the drug.

It is highly improbable that a drug can be developed with the regulatory profile of a drug that exactly matches the genetic (or network) profile of a disease. It is even more improbable to suggest that a drug, designed for another indication, can provide an exact match. Polypharmacology, the design or use of pharmaceutical agents that act on multiple targets or disease pathways, can be utilised to overcome this. Multiple drugs can be combined that each exhibit some overlap with differing subsets of the disease’s regulatory genes. It has been shown that combinatorial drugs exist within the ‘Goldilocks’ zone in gene interaction networks [5]. Within the network, drugs must be sufficiently close to the disease to elicit a change in gene function, but sufficiently distant from the other drugs to ensure there are no antagonistic effects between them. The above study showed that network proximity is correlated with many similarities between drugs, such as gene ontology terms, pathways, clinical, genetic and structural similarity. Ultimately, drugs with lower network proximity will exhibit a larger disparity in terms of mechanism of action. Polypharmacology has seen success in many disease areas such as other cancers and viruses, most notably the human immunodeficiency viruses. We posit that a network polypharmacological approach could provide benefit in identifying treatments for DIPGs (see Figure 1).

**Fig. 1.**
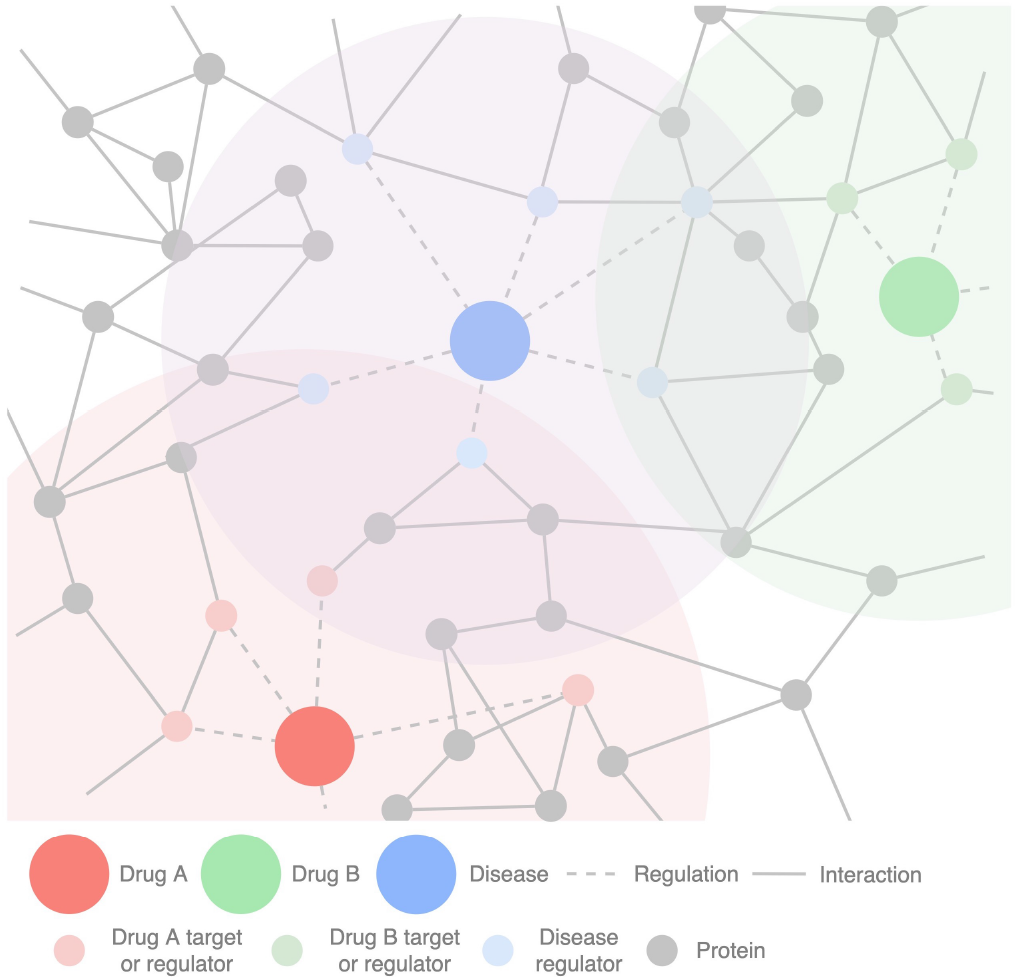
Polypharmacological drug repurposing strategy for DIPG. Illustration displays a tripartite regulatory network of diseases, gene-encoded proteins and drugs. Drugs are shown to have a circle of influence, centered around their drug targets and regulators. This diagram illustrates the ideal drug pair for treatment of DIPG. The two drugs do not share any drug targets or regulators, and their only overlap is genes not thought to regulate the disease.

The interactome (the totality of interactions within a cell), is largely unknown and fascinatingly complex. Drug target interactions constitute much less than one percent of small-molecules reported to bind to a protein. Drug regulatory interactions with disease follow a similar distribution. The lack of known information holds promise that there may be novel indications for current drugs; through the reapplication of drugs to new diseases (through known drug targets), or through novel drug targets.

In recent years, graph-based machine learning (GML) methods have been applied to the task of link prediction, to systemically ‘fill in’ these unreported interactions. Whilst many GML methods exist, the most intuitive is graph convolutional networks. In contrast to conventional neural networks which use arbitrary model architectures, graph convolutional networks models explicitly replicate the network relating to their prediction task. For example, using the aforementioned diseasegene network, and tasked to predict novel gene regulators of disease, genes which are functionally or physically similar are more highly connected within the biological network and by extension more connected in the model architecture. Such genes will have greater influence on each other during the training process of the model. One of the applications of GML is the creation of node embeddings: transforming continuous high dimensional information of node neighbourhoods to low-dimensional dense vectors, to be used in downstream machine learning tasks. Popular approaches include generating embeddings by sampling a network via random walks, treating the sample as corpus of words, and applying natural language processing (NLP) techniques. Similar NLP methods have been applied to amino acid sequences of proteins [6], and molecular substructures of compounds [7] to generate embeddings representing primary structure and molecular substructure of proteins and compounds respectively.

## II. Results

### A. Knowledge Graph

To identify repurposable drugs candidates for DIPG, we must first understand the underlying mechanisms of oncogenesis. To suggest combinatorial drug candidates, we must further understand the biochemical interaction between the therapeutics and assess the synergism of the relationship. From a systems biology point of view, to model this we need to create a network capable of capturing these biochemical and regulatory interactions.

We created a knowledge graph predominantly based on *Pathway Studio*, a biomedical database derived from relationships extracted from over 30 million academic manuscripts (see Methods). Each edge in the graph represents at least one occurrence in scientific literature that has stated a relationship between biological entities (that carboplatin down-regulates DIPG). In total, the graph possessed 1.36 million nodes and 8.52 million edges, weighted according to their frequency in literature (see Fig 2). From this we generated a tripartite subgraph of diseases, drugs and genes. The schema for this subgraph can be seen in Fig 3.

**Fig. 2.**
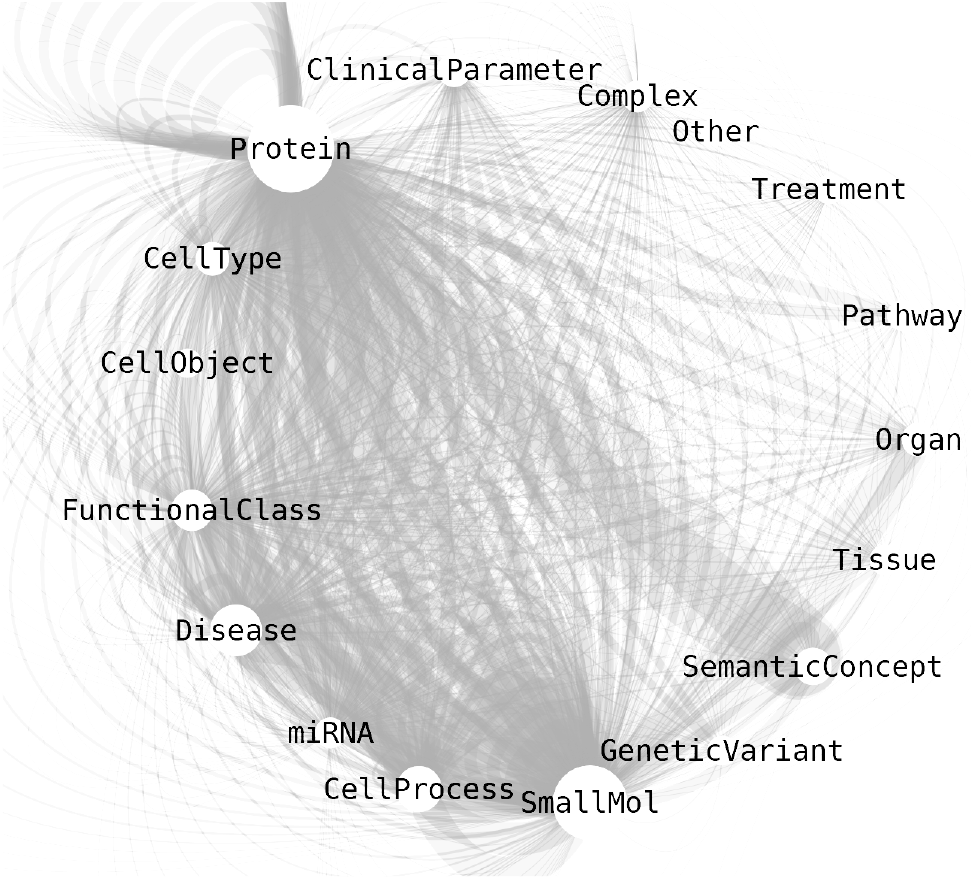
Knowledge graph visualisation using Cytoscape. Size of the node represents the number of nodes of this type in the graph. SmallMol (molecule), Protein and Disease are the largest, alongside SemanticConcept (gene ontology).

**Fig. 3.**
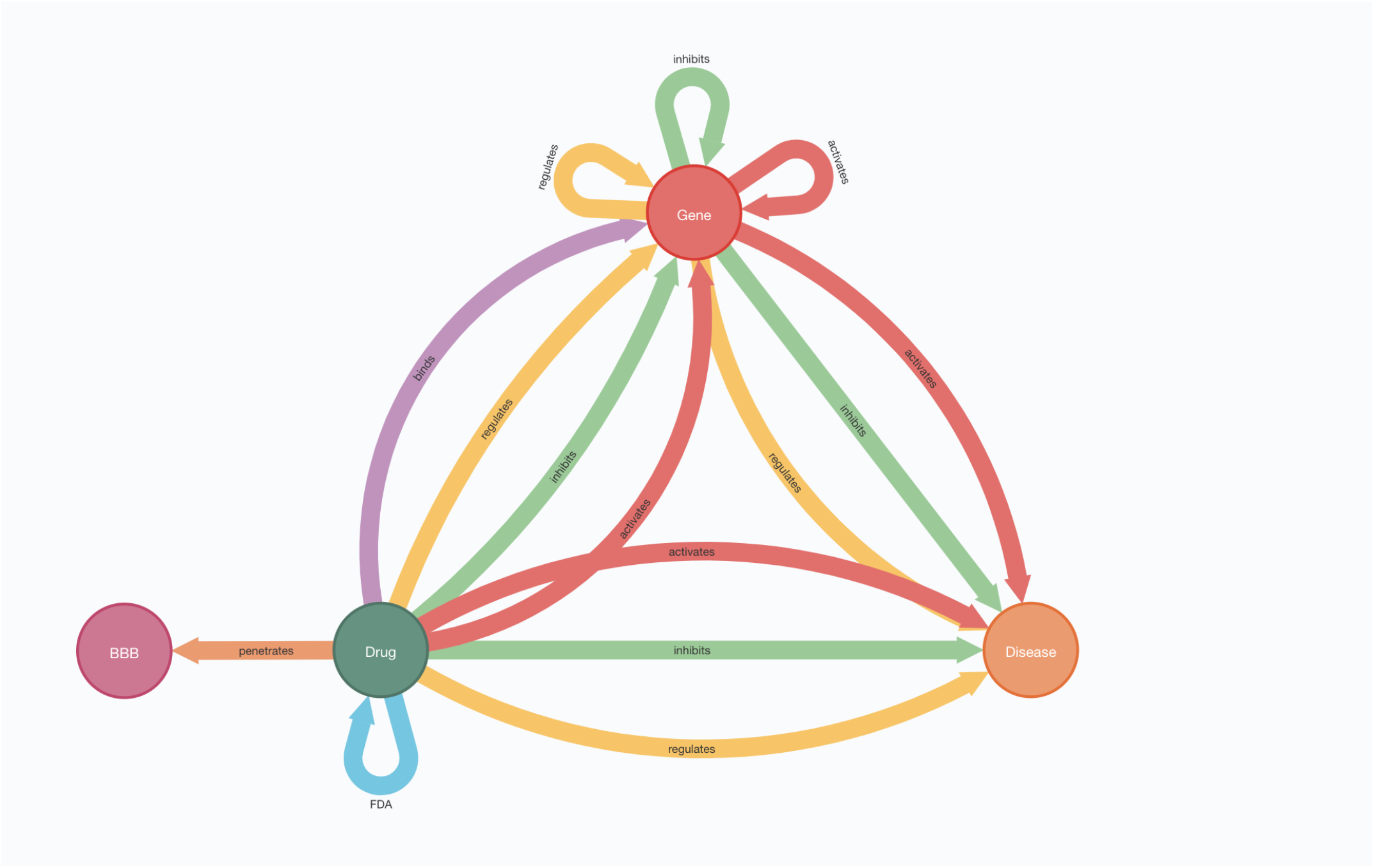
Graph database schema for simplified tripartite graph used in this analysis. Graph consists of drugs, diseases and genes. *Inhibition* and *activation* refer to up- and down-regulation via any direct or indirect mechanism. *Regulation* refers to any direct or indirect mechanism in which the direction of regulation is not known. *Binds* refers to direct binding between drug and gene product. *FDA* refers to a drug pair that are used as part of a multidrug regimen for any indication. *Penetrates* refers to drugs that known to be BBB penetrant.

### B. Blood-brain Barrier Permeability

The success of a drug to treat brain tumors is in part dictated by the compound’s ability to cross the blood-brain barrier (BBB), reaching the tumor at a concentration within the required therapeutic window. To ascertain the BBB permeability of a drug, we trained a link prediction model using embeddings based on molecular sub-structures (see Table 1 I). The selective permeability of drugs crossing the BBB is controlled by: i) ability to passively diffuse through the tight junctions between endothelial cells lining cerebral microvessels, ii) the uptake and efflux transporters and iii) cellular enzyme systems [8]. Our model specifically predicted the ability of a compound to cross the BBB via passive diffusion. We predicted the ability of all approved drugs (approved by either European Medicines Agency or U.S. Food and Drug Administration). The predicted BBB permeability of drugs constituting notable multidrug therapies can be seen in Table V.

**TABLE I.** MODELS USED IN ANALYSIS. *Edge* COLUMN DESCRIBES THE LINK BEING PREDICTED, FROM WHICH ONE OF THE THREE EMBEDDINGS WERE GENERATED. THE *Source* AND *Target* COLUMNS DESCRIBE THE TYPE OF EMBEDDINGS USED FOR THE ADDITIONAL RESPECTIVE SOURCE AND TARGET FEATURES.

| Name | Edge | Source | Target |
| --- | --- | --- | --- |
| CiD | Drug-inhibits-disease | Mol2Vec | Gene-regulates-disease |
| CcC | Drug-combo-drug | Mol2Vec | Mol2Vec |
| CiB | Drug-is-BBB+ | Mol2Vec | - |

### C. Monodrug Inhibition

#### 1) In silico

To ascertain the therapeutic efficacy of drugs for treatment of DIPG, we developed an embedding-based link prediction model capable of predicting the existence of a relationship (otherwise known as an edge) between two nodes. We trained link prediction models to predict drugs that inhibit DIPGs, referred to as the *drug-inhibits-disease* model (see Table I). As this edge is based on occurrence of biological relationships in literature, a predicted link with high probability for the edge *carboplatin-inhibits-DIPG* indicates that if a research group were to research the inhibitory regulation of the drug upon the disease, there is high likelihood the regulation was sufficient to report this regulatory relationship in an academic paper. Top 20 predictions for each disease can be seen in Table For each prediction, we cross-referenced and highlighted known down-regulators reported in literature. From the top 20 predictions, 8 were already known for DIPG (true positive labels in the graph).

#### 2) In vitro

To validate the in silico predictions, we measured cell viability *in vitro*. Using a subset of notable predictions, we used multiple human glioma cell lines derived by surgical biopsy from pediatric H3.3K27M DIPG patients and measured their sensitivity to the drugs over a 72 hour period (see Methods). The 50% inhibitory concentrations (*IC*_50_) for each drug and cell line are tabulated in Table IV. The *IC*_50_ plots of viability as a function of dose can be seen in Figure 4.

**Fig. 4.**
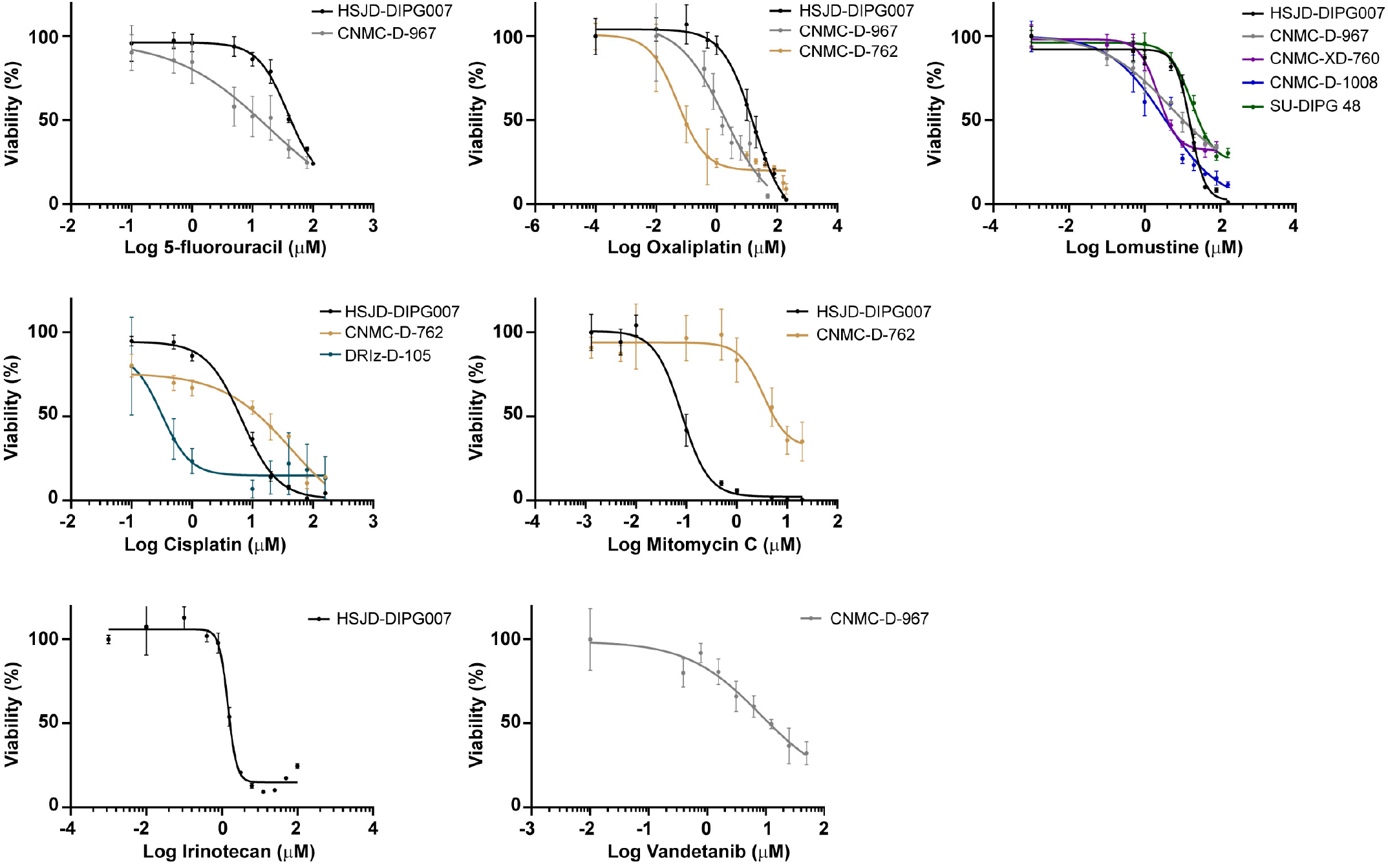
*IC*_50_ measurements of DIPG cell lines treated with anticancer agents. Concentrations were determined using non-linear regression [log (inhibitor) vs. response – variable slope (four parameters)] in GraphPad Prism 8.

### D. Multidrug inhibition

#### 1) In silico

The basis of this proposed drug repurposing methodology is that drug combinations prove more efficacious treatments than compounds administered alone. Having established potential therapeutics, we next wished to predict if a drug pair is complementary. Researchers [5] pioneered a system polypharmacological methodology, demonstrating how using protein-protein interaction networks, network proximity between drug targets of multiple drugs and disease gene regulators can be used to determine the suitability of those two drugs in treatment of the disease. In their work, two proximity measures were employed; distance between drugs (or more precisely distance between the closest drug targets of each drug) and distance between each drug and the disease (distance between the closest drug targets and disease regulators). We posit that by generating node embeddings of a disease-drug regulatory bipartite network, the embeddings will be able to capture both drug-drug proximity and drug-disease proximity. There are notable differences between the original work and ours. Whilst the original metrics represented drug-drug-disease relationships in three dimensions (three one-dimensional proximity measures), we believe that an embedding-based approach would be able to capture more latent information of node neighbourhoods in the higher dimensions of the embeddings. The previous work used protein-protein interactions alongside drug targets and disease regulators to connect drugs and diseases whereas we simply used drug-disease regulation. In a preliminary analysis, we observed extremely high correlation between drug-disease regulation, drug-drug target, and protein-protein interaction bipartite subgraphs (in terms of node embedding similarity and link prediction performance). Hence we believe that, whilst this information is not explicitly supplied to the model, it is implicitly captured in the drug-disease regulatory network alone.

To perform this workflow, we ingested all FDA-approved drug combinations as edges between drugs in our knowledge graph. We assumed that all FDA-approved combinations would exhibit a synergistic or complementary relationship, and thus proved a valuable indicator for the efficacy of drug pairs. We then trained a link-prediction model to predict if drug pairs matched this complementary relationship of FDA-approved drugs using embeddings based on a drug-disease regulatory network and the Morgan substructure of the compounds. Model metrics for the *drug-combo-drug* model can be seen in Table II.

**TABLE II.** MODEL PERFORMANCE OVER 5 RANDOM FOLDS. STANDARD DEVIATION SHOWN. *Name* ABBREVIATION REFERS TO MODELS IN TABLE I.

| Name | AUC | F1 | Accuracy |
| --- | --- | --- | --- |
| CiD | 0.936±0.002 | 0.888±0.003 | 0.888±0.003 |
| CcC | 0.927±0.002 | 0.879±0.001 | 0.876±0.003 |
| CiB | 0.97±0.002 | - | - |

Using the above model, we predicted the likelihood that all the possible pairs of drug candidates (drugs predicted or known to down-regulate DIPG) would exhibit the complementary relationship of FDA-approved drugs. Next, we calculated the transitive probability of combinatorial use of a drug pair to treat DIPG (see Methods). To clarify, the transitive probability of etoposide and doxorubicin for treatment of DIPG, is: the summation of i) the individual probability of inhibition of DIPG from use of etoposide, ii) the individual probability of inhibition of DIPG from use of doxorubicin, and iii) the probability that both drugs are complementary and comply with the relationships of known FDA-approved drug pairs. The top 25 predictions, ranked according to transitive probability for DIPG and are shown in Table V.

#### 2) In vitro

Observing the top 25 multidrug therapies in Table V, we chose 3 notable candidates to test *in silico*. Platinum drugs were by far the highest predicted therapeutics in drug pairs, whilst also showing potent induction of cell death in DIPG cell lines. As such, we decided to choose drugs highly predicted to complement platinum drugs. 5-FU was highly predicted to treat DIPG in combination with both cisplatin and oxaliplatin, which we opted to test. Alongside these drug pairs, we also treated the *CNMC-D-967* cell line with vandetanib in combination with oxaliplatin, similarly highly predicted. All cell lines were treated for 72 hours. Drug interaction landscapes according to the ZIP model are illustrated in Figure 5. All drug combinations were classified as additive. The most synergistic drug pair was 5-FU and oxaliplatin with a ZIP score of 5.71. 5-FU and cisplatin performed the worst, with a ZIP score of 1.75.

**Fig. 5.**
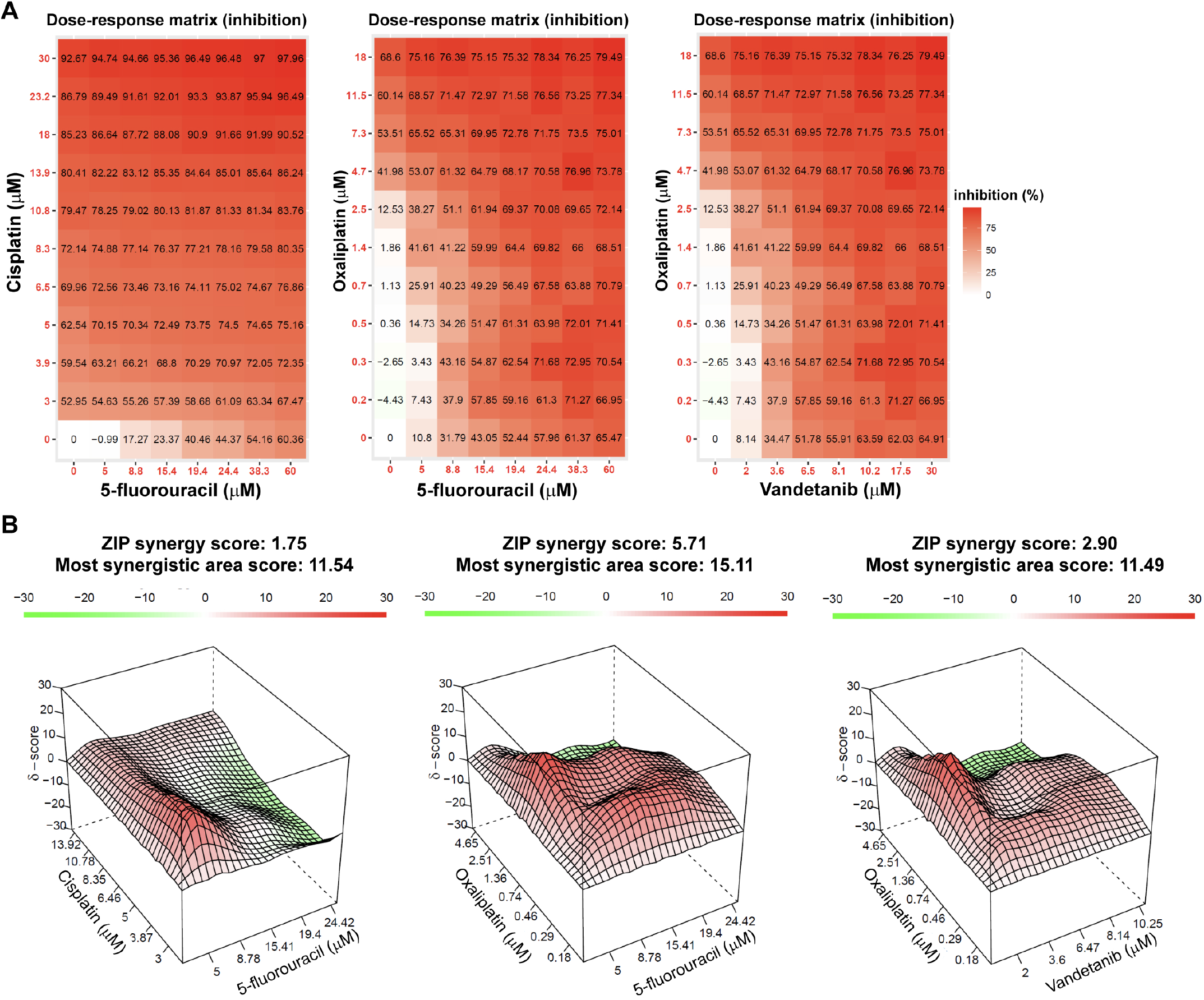
The drug interaction landscape based on the ZIP model. CNMC-D-967 cell line treated with anticancer agents: cisplatin and 5-fluorouracil, oxaliplatin and 5-fluorouracil or oxaliplatin and vandetanib combination for 72 hours. A) 8×11 %Inhibition dose–response matrix. B) calculated 3D synergy maps, SynergyFinder 2.0: visual analytics of multi-drug combination synergies. Synergy score: Less than −10: the interaction between two drugs is likely to be antagonistic; From −10 to 10: the interaction between two drugs is likely to be additive; Larger than 10: the interaction between two drugs is likely to be synergistic.

## III. Discussion

### A. Monodrug Therapies

Using the top predictions of monodrug inhibition of DIPG, we selected notable therapeutics and tested their effect on cytotoxicity *in vitro*. First, we tested 5-fluorouracil (5-FU), a well-established chemotherapeutic for colorectal and other cancers including glioblastoma. The antimetabolite acts through thymidylate synthase inhibition, blocking DNA replication through incorporation of its metabolites into RNA and DNA [9]. 5-FU was the second most highly predicted small molecule inhibitor. Despite the strength of the prediction, 5-FU was poorly effective against each DIPG cell line.

All three platinum containing; cytotoxics, oxaliplatin, cisplatin and carboplatin, were highly predicted to inhibit DIPG. We tested cell viability following treatment of oxaliplatin and cisplatin in multiple cell lines. Interestingly, the *CNMC-D-762* cell line responded to oxaliplatin but not cisplatin, with *IC*_50_values of 0.052 μm and 45.06 μm, respectively. Oxaliplatin is structurally similar to cisplatin, with the exception that two amine groups are replaced by cyclohexyldiamine for improved antitumor activity, and the chlorine ligands are replaced by the oxalato bidentate derived from oxalic acid for improved water solubility. Although such chemical optimizations may explain some of the difference in response, such a disparity in response between platinum analogues is surprising. Oxaliplatin-based regimens have been shown to be superior over cisplatin-based in tumor remission [10]. The physicochemical properties of platinum drugs facilitate relatively good systemic delivery of the drugs, and excellent convection enhanced delivery [11]. Promising CNS penetrance and inhibition suggest oxaliplatin may be a suitable candidate for combinatorial treatment of DIPG.

Lomustine, an alkalating agent, responded well to various cell lines, the lowest of which exhibiting an *IC*_50_ of 2.285 μm (*CNMC-XD-760*). Lomustine is a highly lipophilic nitrosourea compound which undergoes hydrolysis to form reactive metabolites that cause alkylation and cross-linking of DNA and RNA, in turn inducing cytotoxicity. Due to its high lipophilicity, lomustine is highly BBB penetrant and reaches the tumor at therapeutic concentrations. Thus, lomustine may be suitable for use as part of a multidrug regimen.

Next, we tested mitomycin C. The antibiotic functions by selectively inhibiting the synthesis of DNA through the cross-linking of complementary strands. Whilst mitomycin C responded well to the *HSJD-DIPG007* cell line, with an *IC*_50_of 0.081 μm, the drug has been shown to increase levels of P-glycoprotein [12]. P-glycoprotein is one of the major efflux transporters at the BBB, and thus mitomycin may restrict the penetration of various chemotherapeutics.

Irinotecan, a well known topo-isomerase inhibitor, responded well to the *HSJD-DIPG007* cell line with an *IC*_50_ of 1.45 μm. Other topo-isomerases such as etoposide have been used in the treatment of DIPG. Etoposide boasts a higher BBB penetrance and was more strongly predicted to inhibit DIPG, suggesting the drug may be more suitable to treat DIPG than irinotecan.

Overexpression and gene amplification of epidermal growth factor receptor have been reported in a subset of DIPGs [13]. Glioblastomas are some of the most vascularised tumors in which angiogenesis has a critical role in tumorigenesis [14]. Vandetanib, a small-molecule inhibitor of VEGF receptor 2, epidermal growth factor receptor, and RET has been suggested as a potential therapeutic to restore abhorrent levels of EGFR. We tested Vandetanib against the *CNMC-D-967* cell line, however the drug showed only a moderate decrease in induction of cell death, with an *IC*_50_ of 8.138 μm.

### B. Multidrug Therapies

We tested 5-FU with both oxaliplatin and cisplatin, alongside vandetanib with oxaliplatin. Oxaliplatin and 5-FU were highly predicted, and have been suggested as a potential treatment for multiple neoplasms such as advanced colorectal cancer [15]. Despite this, the combination was classified as additive according to the ZIP model of drug synergy. Similarly, cisplatin and 5-FU were classified as additive. Oxaliplatin, the superior analogue of cisplatin showed marginally greater synergy with 5-FU, presumably due the drug’s improvements in terms of antitumor activity and water solubility.

Ultimately, despite all three drug combinations exhibiting high transitive probabilities, their combined effect was not greater than that predicted by their individual potency. Low correlation was found between the *in silico* and *in vitro* models. Whilst all three drug combinations have been proposed as therapies for treatment of DIPG, their relationships were not synergistic. Such a results highlights the unmet need and room for improvement for multidrug therapies for DIPG.

Platinum drugs demonstrated the largest effect on cell viability, therefore one could logically conclude research efforts should focus on finding a supplementary drug that is synergistic with platinum drugs. It is pertinent to remember, however, that platinum drugs demonstrate desirable inhibition of cell growth only in certain cell lines. This variance of response across all cell lines illustrates the heterogeneity of the disease and the complexity of the challenge facing researchers and health-care professionals. It highlights the need for further basic research to stratify such disease endotypes and to understand which endotype respond to which therapy. Until such stratification can be achieved at the point of care, perhaps we should question what kind of relationship a drug pair should ideally possess. (Leaving aside the improbable situation where a wonderdrug or wonder-drug-pair will be developed that inhibits growth of all DIPG endotypes.) Is it desirable to find a synergistic drug combination that inhibits one endotype of DIPG? Or is it more desirable to find an additive drug pair that target two distinct endotypes, to which a greater proportion of patients respond to therapy?

### C. Methodology

One of the main challenges of finding effective treatment of DIPG is the necessity to deliver the drug to the tumor at therapeutic concentrations. Whilst passive diffusion is highly correlated with lipophilicity and thus can be accurately predicted, the situation is much more complex. Efflux transporter proteins (such as those belonging to the adenosine triphosphate binding cassette superfamily), have a major role in the delivery of chemotherapeutics. It is becoming increasingly clear many of the common drugs used to treat glioblastomas have an effect on efflux transporters such as P-glycoprotein, the multidrug resistance protein or the breast cancer resistance protein. Whilst our analysis highlighted drugs predicted to be BBB penetrant via passive diffusion, it did not take into consideration interactions between the potential drugs and efflux transporters.

The *drug-combo-drug* pairwise model finds high dimensional correlations between drugs (in terms of node neighbourhood and physicochemical properties of the compounds) to determine if two drugs are suitable for combinatorial use (for any indication). Due to the unique requirement of brain cancers to be BBB penetrant, the model would not take this into consideration. Models are only as good as the data they are trained on. As there is no gold standard therapeutic to treat glioblastomas, the *drug-inhibits-disease* model was trained on less than ideal positives (regulators reported in literature to inhibit brain neoplasms but have poor BBB penetrance), and thus predictions will reflect this obfuscation.

*In silico* and *in vitro* disease models are approximations of a disease. Such approximations have their shortcomings. In terms of our computational analysis, an important note is that the above drugs have vastly differing degrees of connectivity in the knowledge graph. A well-used drug such as imatinib is used in the treatment of many neoplasms and thus has many reported regulation edges in the graph. Lesser known kinase inhibitors such as trametinib have an order of magnitude fewer edges. Such differences in degree indicate different prior probabilities of treatment. In other words, nodes in a graph with higher connectivity are statistically more likely to be connected. This prior probability is reflected in the probability distributions. Imatinib will have a considerably higher average probability compared to trametinib thus the model will be biased to predict more highly connected drugs. All of the top 20 predicted inhibitors of DIPG are well-known therapeutics for treatment of the disease, indicating the prevalence of this bias. In contrast, high throughput screening of multiple DMG cultures (without this bias) yielded promising investigational drugs such as marizomib [16]. It is difficult to ascertain to what extent predictions are dependent on the therapeutic efficacy of the drug, or simply on this literature bias. Recently researchers have attempted to prevent the encoding of node degree into embeddings [17], [18]. It would be interesting to see if the use of such embeddings in a link prediction model would highlight more unknown left-of-field therapeutic candidates.

For our *in silico* models, we used a literature-derived knowledge graph. The DIPG disease node represents any occurrence in literature that referenced the disease. To clarify, the disease node is a conflation of all *in vitro* and *in vivo* models, and any clinical observations. All disease endotypes are similarly conflated. There is a known lack of agreement between observations seen in *in vitro* DIPG models and *in vivo*, thought to be primarily due to the mechanisms preventing drug delivery. Whilst a machine learning model that predicted *IC*_50_ values for a specific cell line would be more correlated with the experimentally validated cell viability, this would ignore the problem of drug delivery. By using a literature-derived method (that involves a conflation of all methods) we hoped this would provide more less biased results. One obvious shortcoming of this methodology is there is no differentiation between disease subtypes. As can be seen in Table IV, drug responses differ greatly between different patient-derived cells. Combining endotypes with obviously disparate cellular environments may be naïve and impede understanding of the underlying tumorigenic mechanisms.

**TABLE III.**
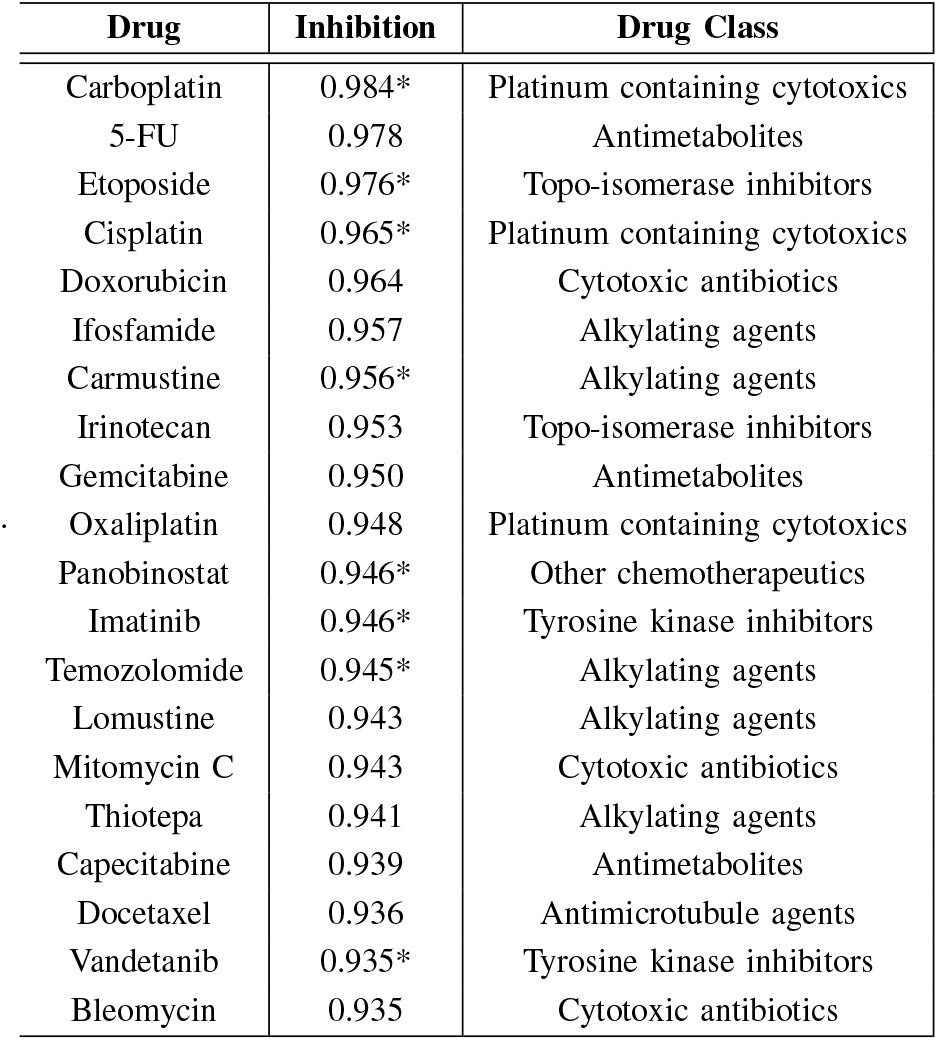
TOP 20 PREDICTED INHIBITORS OF DIPG. * INDICATES DISEASE REGULATION IS ALREADY KNOWN

**TABLE IV.**
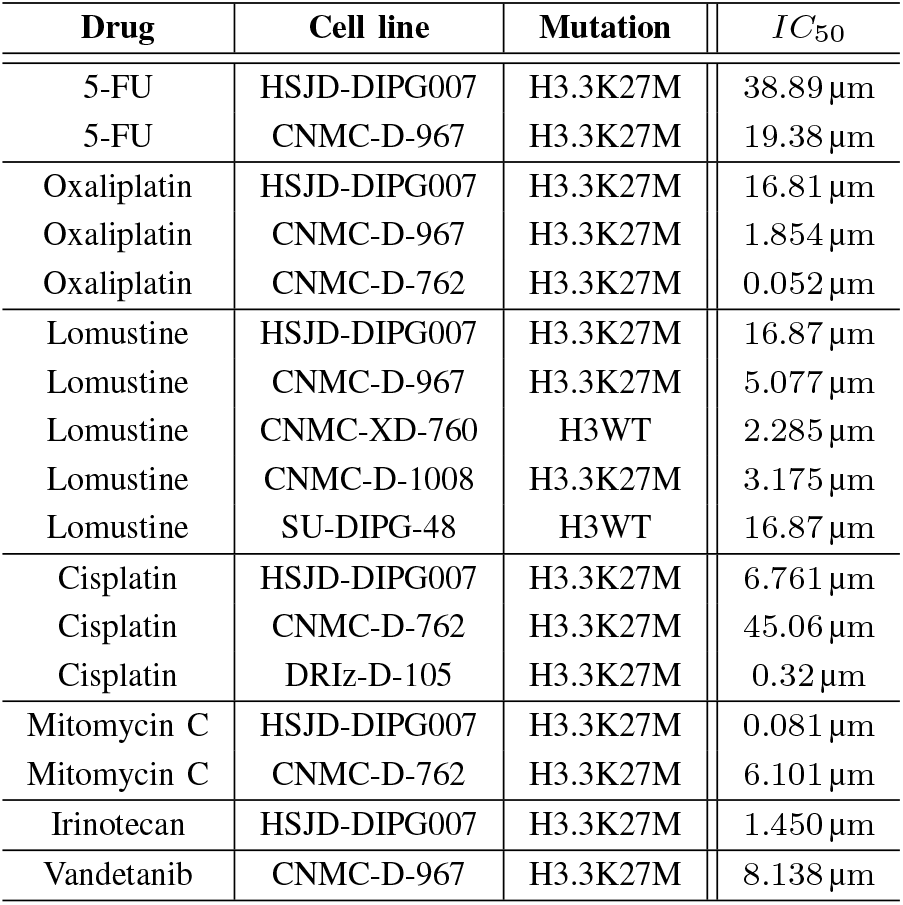
*IC*_50_ VALUES AT 72 HOURS AFTER TREATMENT WITH ANTICANCER AGENTS IN DIPG CELL LINES

| Drug | Cell line | Mutation | $IC_{50}$ |
| --- | --- | --- | --- |
| 5-FU | HSJD-DIPG007 | H3.3K27M | 38.89 μm |
| 5-FU | CNMC-D-967 | H3.3K27M | 19.38 μm |
| Oxaliplatin | HSJD-DIPG007 | H3.3K27M | 16.81 μm |
| Oxaliplatin | CNMC-D-967 | H3.3K27M | 1.854 μm |
| Oxaliplatin | CNMC-D-762 | H3.3K27M | 0.052 μm |
| Lomustine | HSJD-DIPG007 | H3.3K27M | 16.87 μm |
| Lomustine | CNMC-D-967 | H3.3K27M | 5.077 μm |
| Lomustine | CNMC-XD-760 | H3WT | 2.285 μm |
| Lomustine | CNMC-D-1008 | H3.3K27M | 3.175 μm |
| Lomustine | SU-DIPG-48 | H3WT | 16.87 μm |
| Cisplatin | HSJD-DIPG007 | H3.3K27M | 6.761 μm |
| Cisplatin | CNMC-D-762 | H3.3K27M | 45.06 μm |
| Cisplatin | DRIZ-D-105 | H3.3K27M | 0.32 μm |
| Mitomycin C | HSJD-DIPG007 | H3.3K27M | 0.081 μm |
| Mitomycin C | CNMC-D-762 | H3.3K27M | 6.101 μm |
| Irinotecan | HSJD-DIPG007 | H3.3K27M | 1.450 μm |
| Vandetanib | CNMC-D-967 | H3.3K27M | 8.138 μm |

**TABLE V.** TOP 25 PREDICTED DRUG COMBINATIONS FOR DIPG. * INDICATES DISEASE REGULATION ALREADY KNOWN, ^†^ INDICATES DRUG PAIR ALREADY FDA-APPROVED FOR ANOTHER INDICATION, ^‡^ INDICATES DRUG IS KNOWN TO PERMEATE THE BBB.

| Drug 1 (D1) | Drug 2 (D2) | Inhibit D1 | Inhibit D2 | Combination | BBB+ D1 | BBB+ D2 | Transitive |
| --- | --- | --- | --- | --- | --- | --- | --- |
| Carboplatin | Cisplatin | 0.984* | 0.965* | 9.411<sup>†</sup> | 0.833 | 0.958 | **0.714** |
| Cisplatin | Temozolomide | 0.965* | 0.945* | 0.882<sup>†</sup> | 0.958 | 0.920 | **0.710** |
| Cisplatin | Ifosfamide | 0.965* | 0.957 | 0.912<sup>†</sup> | 0.958 | 0.872 | **0.704** |
| Cisplatin | Oxaplatin | 0.965* | 0.948 | 0.929<sup>†</sup> | 0.958 | 0.794 | **0.657** |
| Cisplatin | Cyclophosphamide | 0.965* | 0.847 | 0.942<sup>†</sup> | 0.958 | 0.829 | **0.611** |
| Carboplatin | Oxaplatin | 0.984* | 0.948 | 0.960<sup>†</sup> | 0.833 | 0.794 | **0.593** |
| Cisplatin | Imatinib | 0.965* | 0.946* | 0.904<sup>†</sup> | 0.958 | 0.746 | **0.590** |
| Carboplatin | 5-FU | 0.984* | 0.978 | 0.962<sup>†</sup> | 0.833 | 0.757 | **0.584** |
| 5-FU | Cisplatin | 0.978 | 0.965* | 0.815<sup>†</sup> | 0.757 | 0.958 | **0.558** |
| Cisplatin | Thalidomide | 0.965* | 0.917* | 0.685 | 0.958 | 0.956<sup>‡</sup> | **0.555** |
| Carboplatin | Temozolomide | 0.984* | 0.945* | 0.766 | 0.833 | 0.920 | **0.546** |
| Carboplatin | Ifosfamide | 0.984* | 0.957 | 0.795 | 0.833 | 0.872 | **0.545** |
| Cisplatin | Gemcitabine | 0.965* | 0.950 | 0.872<sup>†</sup> | 0.958 | 0.705 | **0.540** |
| 5-FU | Oxaliplatin | 0.978 | 0.948 | 0.929<sup>†</sup> | 0.757 | 0.794 | **0.518** |
| Cisplatin | Lenalidomide | 0.965* | 0.825 | 0.737 | 0.958 | 0.921 | **0.517** |
| Cisplatin | Tegafur | 0.965* | 0.773 | 0.926<sup>†</sup> | 0.958 | 0.780 | **0.516** |
| Cisplatin | Gefitinib | 0.965* | 0.870 | 0.901<sup>†</sup> | 0.958 | 0.706 | **0.512** |
| Carboplatin | Thalidomide | 0.984* | 0.917* | 0.712 | 0.833 | 0.956<sup>‡</sup> | **0.512** |
| Ifosfamide | Oxaliplatin | 0.956 | 0.948 | 0.808 | 0.872 | 0.794 | **0.508** |
| Temozolomide | Vorinostat | 0.945* | 0.805 | 0.847<sup>†</sup> | 0.920 | 0.849 | **0.503** |
| Cyclophosphamide | Lenalidomide | 0.847 | 0.825 | 0.936<sup>†</sup> | 0.829 | 0.921 | **0.499** |
| Oxaliplatin | Vandetanib | 0.948 | 0.935* | 0.879<sup>†</sup> | 0.794 | 0.804 | **0.497** |
| Thalidomide | Cyclophosphamide | 0.917* | 0.847 | 0.807<sup>†</sup> | 0.956<sup>‡</sup> | 0.829 | **0.497** |
| 5-FU | Ifosfamide | 0.978 | 0.957 | 0.802 | 0.757 | 0.872 | **0.496** |
| Carboplatin | Gemcitabine | 0.984* | 0.950 | 0.901<sup>†</sup> | 0.833 | 0.705 | **0.495** |

Link prediction on knowledge graphs signifies that the area under the receiver operating characteristic curve is not 0.5 for random guesses of connecting source and target nodes: it is much higher. Researchers have shown prior probability of treatment (the likelihood two nodes are connected simply by their node degrees) can be achieved by randomly permuting bipartite graphs multiple times (swapping edges but preserving node degree) and noting the average probability of an edge as a function of source and target degree. Ensuring an edge prediction is larger than the prior probability of connection, guarantees local network topology suggests the presence of an unreported edge, not simply the global degree distribution. The prior probability dominates predictions on most networks. One recent attempt to overcome this uses a Bayesian approach during the training process of the graph embeddings; modelling the prior probability as the prior in the Bayes formula [18]. By explicitly modelling the prior, it ensures it is not captured in the graph embedding.

Machine learning methods are often cursed by their lack of explainability. Specifically, the optimized parameters of supervised models are hard to interpret. Whilst embeddings methods have proved powerful strategies to capture continuous data into discrete and dense vector representations, they obfuscate explainability even further. Embeddings are used as the features of a model. As each dimension is the output of another vectorization model, they are completely uninterpretable.

FDA approved drugs were used as a proxy to measure the complementary action of two drugs. This may not be the best approach to ascertain this relationship. Drug synergy dramatically varies between cell lines. If one wished to predict synergy for certain drugs pairs in certain cell lines, there are more established and accurate methods available [19]. Whilst FDA approval was shown to be correlated with drug synergy, the majority of drug pairs are different drug classes such as antivirals and kinase inhibitors are often indicated together. If one wished to predict complementary drugs acting upon different disease mechanisms, it may be beneficial to remove all drug pairs from the same drug class from the training set of the model. Alternatively, this could easily be achieved by imposing a threshold of network proximity (cosine distance), to ensure functional and mechanistic dissimilarity between drugs.

This workflow trained and predicted on only FDA-approved drugs. Whilst there are a few thousand drugs in the graph, there are over 1 million compounds. Rerunning the analysis on all compounds may highlight i) drug-like compounds with therapeutic potential and ii) potential nutraceutical candidates. We did not initially include compounds, as the degree of connectivity massively varies between drugs and compounds, to a much greater extent than diseases. We believed this increased prior probability of connection would have dominated the predictions. The prior probability issue should be resolved before rerunning the analysis.

Biomedical knowledge graphs are largely incomplete. On average, well under one percent of possible relationships are reported for edges such as regulation, binding and expression. This presents a problem for link prediction models; the true distribution of positive and negative classes is unknown, and it is impossible to differentiate a true (but unreported) positive from a false positive. It is preferable to train your model on a class balance that matches your real world distribution. In this application, this is not possible. Moreover, when your real world distribution is known, optimizing your classification threshold becomes a much simpler task. We used maximal f1 score to determine the optimal threshold. This often meant an unrealistic increase in the number of positive predicted samples. For 3302 drugs and 19387 diseases, there are 163590 known inhibitors (0.25 percent). The average positive class for a random subset of predicted regulators above the optimal threshold of 0.634 was 28 percent, a 112-fold increase in regulators. This is obviously a gross overestimation in the number of unreported regulators and a major limitation in our approach. We encourage predictions to be analysed only in the context of the relative list of regulators for that disease. An alternative approach taken to decide classification threshold would be based largely on the known class distribution [4].

The knowledge graph provides an incredible amount of biological information, and we only used a small amount of this information. Each edge in the graph was weighted according to the absolute occurrence that this relationship appears in scientific literature. It is logical to believe the number of times a relationship is reported is correlated with its strength or confidence. This does, however, introduce a new level of knowledge bias. Genes such as p53 have received tremendous attention. Does this signify said gene’s importance over it’s less researched genetic counterparts? In previous unpublished analyses, we attempted to utilise this weighting through numerous normalisation strategies such as frequency of node, edge and path and log-scaled reference counts. We found that such strategies only generalised solutions; wherein well-researched areas performed better, whilst less-researched performed worse. An interesting approach would be to normalise nodes according to the weighted edges that connect to it (weighted by reference count). In this case, for a well-researched node with 10 edges and an individual edge weight of 50, each edge would be equivalent to a node with 10 edges and an edge weight of 5. This may also solve the prior probability issue stated earlier, as for nodes with fewer edges, each edge will be given higher importance, partially mitigating the dominating effect of nodes with high degrees.

## Conclusion

There are currently no satisfactory means of treatment of DIPG. Using a network polypharmacology approach, we highlighted multidrug therapeutics to be used for the treatment of DIPG. We calculated transitive probabilities for each drug pair based on the predicted regulatory action and blood-brain barrier permeability of each drug, and the predicted synergy of the drug pair. Whilst top predictions yielded drug combinations commonly used for other neoplasms, the drug pairs were seen to be only additive when measured in a combinatorial drug synergy assay. DIPG cell lines showed dramatically different responses when treated by monodrug therapies, illustrating the heterogeneity of the disease. Challenges such as this heterogeneity, difficulties of modelling brain permeability, and an inherent bias in literature-derived link prediction methods need to be overcome before a satisfactory treatment for DIPG is developed.

## IV. Methods

### A. Knowledge Graph

We created a knowledge graph predominantly based on Pathway Studio, a literature-derived database that uses natural language processing techniques to leverage biological relationships from over 30 million literary sources. Pathway Studio also contains the relevant subset of Reaxys Medicinal Chemistry, a database of small-molecule protein bioactivities, pertaining the species homo sapiens, Mus muccus, and Rattus rattus, and rattus norvegicus. We appended this core graph with gene ontologies [20], drug side effects from SIDER [21] and drug-target information from Drug Central [22]. We also created similarity links such as protein-protein similarity (Local Smith Waterman of over 0.5), and molecule substructure similarity (Tanimoto similarity of Morgan Fingerprint of over 0.5). After refactoring and harmonization, the graph possessed 1.36 million nodes and 8.52 million edges and over 200 edge types, weighted according to their occurrences in literature.

For each molecule with an InChI code or key within the graph, we generated a Mol2Vec embedding, using the pretrained embeddings [7] were trained on 20 million compounds from the ZINC database [23]. As all other embeddings had a dimension of 100, and as the model requires embeddings of the same size, we used the scikit-learn version of principal component analysis to reduce the embedding down to the required size. Summation of the explained variance showed only 1.3 percent of information was lost. Similarly, we generated embeddings for proteins based on trimers of their amino acid sequence using the pretrained model of ProtVec [6], trained on 551,754 proteins from Swiss-Prot. Because trimers can start at the first, second or third amino acid in a protein sequence, three embeddings were generated per protein. As per the methodology of the original paper, we took the element-wise average of these.

For this analysis, we created a tripartite subgraph of diseases, genes and drugs, connected via multiple regulatory edges. Said edges included up- and down-regulation via any direct or indirect mechanism. For example, the edge *drug-inhibits-gene* conflates expression, indirect regulation, direct binding (agonism and antagonism), and promoter binding. We also provided a conflated edge of both up- and down-regulation.

### B. Monodrug Inhibition Prediction

To determine if a drug down-regulated a disease or gene, we developed an embedding-based link prediction model based on multiple disease regulatory bipartite networks and additional physicochemical and structural information of the source and target nodes (see Fig 6). The embeddings were used by a random forest classifier (scikit-learn implementation), optimized via hyper-parameter bayesian optimization. We assessed multiple node embedding strategies in this project. For embedding choice in the link prediction model, we assessed GraRep, nodevec, LINE and SVD model. All models used the BioNev [24] implementation, except node2vec, for which we used the C++ version from SNAP [25]. Random search was employed to determine models with the highest AUC score. We also investigated different mathematical functions to create an edge embedding from two node embeddings (concatenation, element-wise average, hadamard, L1 and and L2 loss). Because our model used two different types of embeddings for each edge (for example: i) graph embedding for protein and molecule, and ii) ProtVec and Mol2vec respectively), employing the hadamard edge function signifies, the hadamard was calculated both for graph embeddings, and the for Mol2vec and ProtVec separately before concatenating. We also investigated stacking models to create soft-voting bagging classifiers. Because a subset of edges must be removed and used to train the model, we postulated that training multiple models on different subsets, and combining their predictions via a weighted average according to the performance, would increase predictive power of the stacked model. Results showed, however, that increase was extremely marginal (under 0.5 percent increase in AUC). Due to the doubling of training time, we deemed this increase unnecessary. The following metrics were used to determine model performance: AUC, F1 score, accuracy, and precision. The best classification threshold was determined by the finding at which F1 score was highest (the point at which accuracy and precision intersect). Scores were averaged over 5 random folds.

**Fig. 6.**
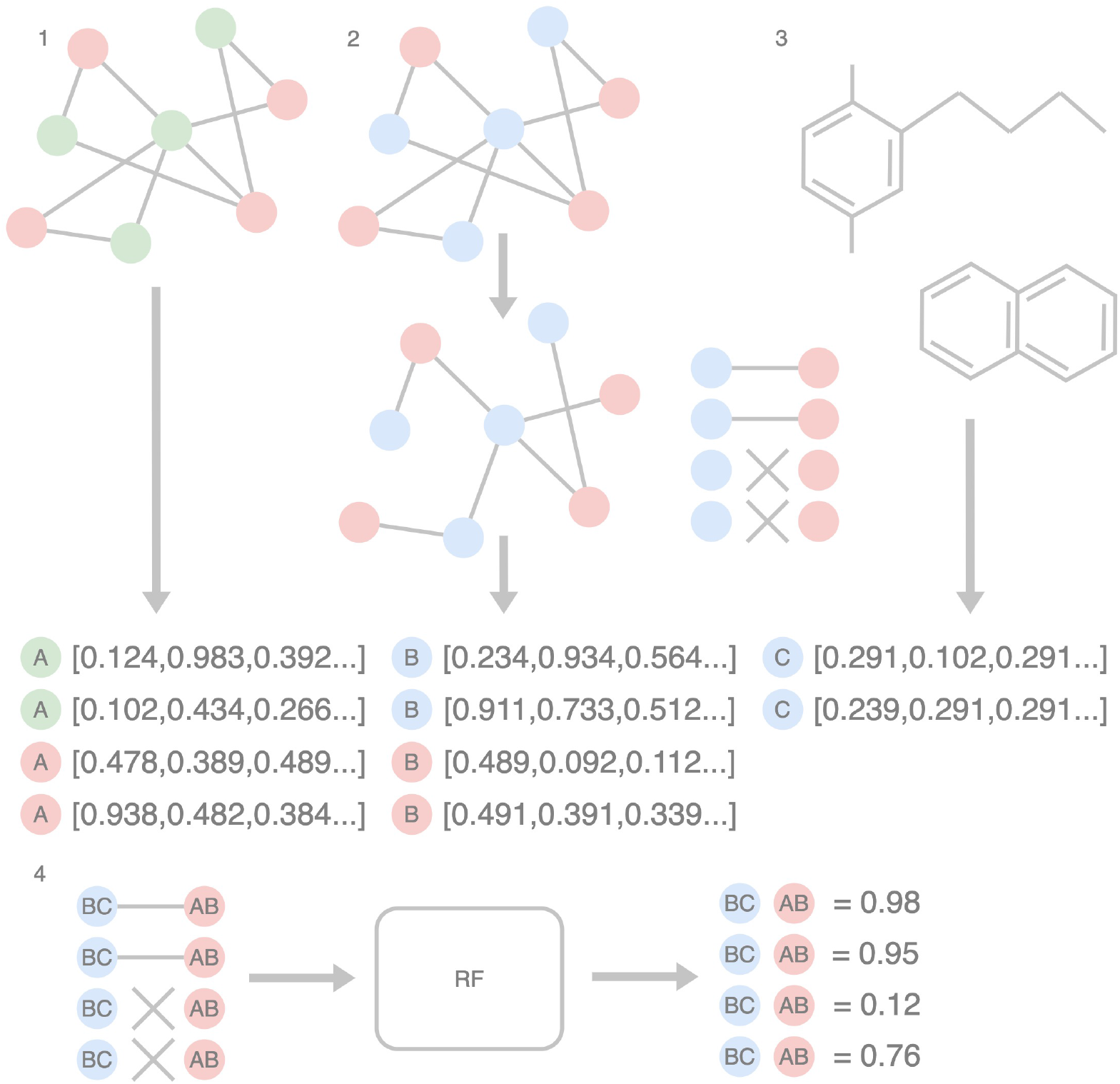
Workflow for model generation for *drug-inhibits-disease* edge. First, we split the graph into two subgraphs, a disease-gene subgraph (Red and green nodes, respectively), and a disease-mol subgraph (red and blue, respectively). 1) We generated embeddings for disease and gene nodes (Embeddings A). 2) We removed a subset of the edges of the disease-mol subgraph and generate node embeddings for disease and molecules (Embeddings B). 3) We generated embeddings based on the molecular substructure (Embeddings C). We now have two embeddings for each disease, and two for each molecule. To summarise, Embeddings A describe the neighbourhood or proximity of disease and genes in a disease-gene regulatory network (how close is one disease to another). Embeddings B describes the neighbourhood or proximity of a disease and molecule in a disease-mol regulatory network. Embeddings C describe the physicochemical similarity of compounds. 4) We combined the two embeddings for each disease and molecule (AB and BC, respectively), and used the edges removed from the disease-mol network to train a model capable of predicting the existence of a link between a disease and a molecule. In the example below, we can see there were two known disease-gene pairs (the regulation has been stated in Pathway Studio), and two random disease-gene pairs. We can see that the model has predicted the final pair to actually be an unreported regulation link.

For the protein-drug regulation prediction, node neighbourhood embeddings of dimension *d* for source and target *n*_*s,t*_, were complemented with structural information: *e*_*s*_ protein ProtVec amino acid trimer embeddings, and *e*_*t*_ Mol2Vec Morgan fingerprint substructure embeddings, where:

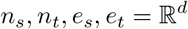

The L2 norm between pairs of node and structural embeddings were calculated via L2 before the vectors were concatenated together:

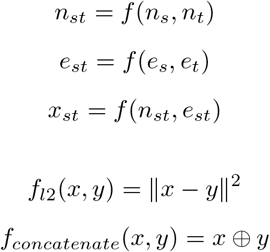

### C. Drug Combination Prediction

We ingested FDA approved drugs from the DrugCombDB dataset [26], mapped to the drugs in our knowledge graph by PubChem CID. As multidrug therapies exist within the dataset (more than two drugs), we split these into all possible two pair combinations of these multidrug combinations, and individually added these to the graph. Next, we trained a link-prediction model to predict the FDA Combination relationship between two drugs. This model differed from the previous models used to predict down-regulation of diseases, genes and cellular processes. The previous models generated embeddings of the bipartite subgraph of the edge it was predicting, alongside an additional embedding based on an additional network. To clarify, the model to predict *drug-inhibits-disease* edges used embeddings based on a *drug-inhibits-disease* subgraph alongside embeddings based on a *gene-inhibits-disease* subgraph. In contrast, this model, trained to predict the *drug-combo-drug* edge, did not generate embeddings from a *drug-combo-drug* bipartite graph, whereas it used only a *drug-combo-drug* subgraph. As the GraRep embedding is transductive, one can only generate embeddings if the node exists within the graph. Thus, using embeddings based on *drug-combo-drug* would limit predictions to the 572 drugs currently approved for combinatorial use. In contrast, using the *drug-combo-drug* subgraph permitted predictions for 2861 drugs. We investigated the effect of using the FDA combination edges alongside the drug-disease regulatory edges, which yielded a 2 percent increase in AUC. We deemed the decrease in utility and drug scope more important than this increase in model performance.

We calculated transitive probability of treatment 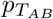 using the equation below, where 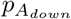 is the probability drug A down-regulates the disease, 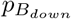 is the probability drug B down-regulates the disease and 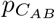 is the probability drug A and drug B are FDA-approved (complementary):

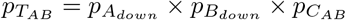

### D. Blood-brain Barrier Permeability Prediction

To predict if a drug was BBB+ or BBB−, we created a simple predictive model. For training data, we used Adenot and Lahana’s dataset [27]. Once mapped onto molecules in our graph, there were 1220 BBB+ drugs and 291 BBB-. This class imbalance threatened to dominate the predictions towards the majority class (BBB+). To prevent this, we used Synthetic Minority Oversampling Technique (SMOTE) to artificially synthesise negative samples. We used pretrained Mol2Vec embeddings, based on Morgan substructures, trained on 20 million molecules from ZINC [23] as features for our model. The pretrained Mol2Vec embeddings had a dimension of 300. Since the minority class had only 291, we applied principal component analysis (PCA) to reduce the embeddings until 0.95 of the variance was still conserved. This created reduced embeddings of 35 dimensions; a significant reduction in vector size, whilst only losing 0.05 percent of information within the embeddings. We used a Bayesian-optimized random forest classifier for this model. To validate the model we used 10-fold stratified cross validation. Importantly, to prevent data leakage, SMOTE and PCA were applied within each fold of the training data. Artificially generated samples were only used for training and not testing.

A previous study [28] showed drug indications and side effects can be used to predict if a drug is BBB+ or BBB-. Their logic dictates that any drug that exhibits clinical phenotypes associated with the brain must be able to pass or circumvent the BBB via any mechanism. The study used grouped brain-associated phenotypes as features. We replicated this analysis by replacing Mol2Vec with embeddings based on i) drug-disease regulatory network and ii) drug-disease association network. We reasoned that the embeddings would be able to capture the brain-associated diseases and therefore would be informative to the model. The replacement with both of these embeddings represented an 5 percent decrease in AUC score.

We calculated the transitive probability of treatment 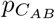 using a similar equation as in Drug Combinations, however extended to include the probability of BBB permeability for each drug (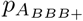 and 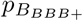 for drug A and drug B respectively).

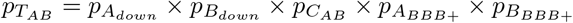

### E. Cell Viability

#### 1) Patient-derived cell cultures

All human cell cultures were generated from biopsy or autopsy samples collected in accordance with informed consent and in compliance with the dissociation protocol for DIPG cells from Biopsies established at the Children’s National Hospital (CNMC) in Washington (IRB protocols, #1339) and DIPG Research Institute Zurich (BASEC-Nr.: 2019-00615). HSJD-DIPG007 cells were kindly provided by Dr. Angel Montero Carcaboso at the Hospital Sant Joan de Deu, Barcelona. Cells were maintained in NeuroCult NS-A Basal Medium with NS-A Proliferation Supplement (STEMCELL Technologies, Vancouver, CA), 1X Antibiotic/Antimycotic (ThermoFisher), 40 ng/mL epidermal growth factor (PeproTech, NJ, USA), and 40 ng/mL fibroblast growth factor (PeproTech, NJ, USA). All cell culture models were validated by DNA fingerprinting.

#### 2) Drugs and Cell Viability Assays

Drugs used in this body of work were purchased from Selleckchem. For drug treatment cell viability cells were plated in 96-well plates at 5000 cells per well and cultured 72 hours in the presence of drug in quadruplicate. Experiments were repeated for validation. Cell viability was measured using a CellTiter-Glo R Luminescent Cell Viability Assay (G7570, Promega) and data were collected on a Biotek Cytation 3 luminescence reader. *IC*_50_ concentrations were determined using non-linear regression [log (inhibitor) vs. response – Variable slope (four parameters)] in GraphPad Prism 8 (LaJolla, CA).

#### 3) Combinatorial Drug Synergy

For testing combinatory effects of two drugs, the cells were treated with each drug individually or in combination for 72 hours before subjecting to CellTiter-Glo assay. Drug interactions were evaluated using the SynergyFinder 2.0 platform [29], which uses the Zero Interaction Potency (ZIP) [30] model for quantifying drug synergy.

## V. ACKNOWLEDGMENT

This project was a *pro bono* collaborative project between The Anticancer Fund, University Children’s Hospital Zurich, and Elsevier. The authors would like to express their gratitude to all parties for affording the resource to undertake this project. The analysis used the tremendously useful Pathway Studio, to which we thank Anton Yuryev.

## REFERENCES

[1] P. Pantziarka, C. Verbaanderd, I. Huys, G. Bouche, and L. Meheus, “Repurposing drugs in oncology: From candidate selection to clinical adoption,” in Seminars in Cancer Biology. Elsevier, 2020.

[2] “Existing drug may treat the deadliest childhood brain tumor, stanford-led study finds.” [Online]. Available: https://www.stanfordchildrens.org/en/about/news/releases/2015/existing-drug-may-treat-the-deadliest-childhood-brain-tumor

[3] D. S. Himmelstein, A. Lizee, C. Hessler, L. Brueggeman, S. L. Chen, D. Hadley, A. Green, P. Khankhanian, and S. E. Baranzini, “Systematic integration of biomedical knowledge prioritizes drugs for repurposing,” eLife, vol. 6, 2017.

[4] F. Womack, J. Mcclelland, and D. Koslicki, “Leveraging distributed biomedical knowledge sources to discover novel uses for known drugs,” Nov 2019.

[5] F. Cheng, K. I. A., and B. Albert-Laszlo, “Network-based prediction of drug combinations,” Nature Communications, vol. 10, no. 1, 2019.

[6] E. Asgari and M. R. K. Mofrad, “Continuous distributed representation of biological sequences for deep proteomics and genomics,” Plos One, vol. 10, no. 11, Oct 2015.

[7] S. Jaeger, S. Fulle, and S. Turk, “Mol2vec: Unsupervised machine learning approach with chemical intuition,” Journal of Chemical Information and Modeling, vol. 58, no. 1, p. 27–35, Oct 2018.

[8] H. Sherman and A. E. Rossi, “A novel three-dimensional glioma blood-brain barrier model for high-throughput testing of tumoricidal capability,” Frontiers in oncology, vol. 9, p. 351, 2019.

[9] D. B. Longley, D. P. Harkin, and P. G. Johnston, “5-fluorouracil: mechanisms of action and clinical strategies,” Nature reviews cancer, vol. 3, no. 5, pp. 330–338, 2003.

[10] F. Zhang, Y. Zhang, Z. Jia, H. Wu, and K. Gu, “Oxaliplatin-based regimen is superior to cisplatin-based regimen in tumour remission as first-line chemotherapy for advanced gastric cancer: A meta-analysis,” Journal of Cancer, vol. 10, no. 8, p. 1923, 2019.

[11] F. E. El-Khouly, D. G. van Vuurden, T. Stroink, E. Hulleman, G. J. Kaspers, N. H. Hendrikse, and S. E. Veldhuijzen van Zanten, “Effective drug delivery in diffuse intrinsic pontine glioma: a theoretical model to identify potential candidates,” Frontiers in Oncology, vol. 7, p. 254, 2017.

[12] A. Noack, S. Noack, A. Hoffmann, K. Maalouf, M. Buettner, P.-O. Couraud, I. A. Romero, B. Weksler, D. Alms, K. Römermann et al., “Drug-induced trafficking of p-glycoprotein in human brain capillary endothelial cells as demonstrated by exposure to mitomycin c,” PLoS One, vol. 9, no. 2, p. e88154, 2014.

[13] R. J. Gilbertson, D. A. Hill, R. Hernan, M. Kocak, R. Geyer, J. Olson, A. Gajjar, L. Rush, R. L. Hamilton, S. D. Finkelstein et al., “Erbb1 is amplified and overexpressed in high-grade diffusely infiltrative pediatric brain stem glioma,” Clinical cancer research, vol. 9, no. 10, pp. 3620–3624, 2003.

[14] A. Broniscer, J. N. Baker, M. Tagen, A. Onar-Thomas, R. J. Gilbertson, A. M. Davidoff, A. P. Panandiker, W. Leung, T. K. Chin, C. F. Stewart et al., “Phase i study of vandetanib during and after radiotherapy in children with diffuse intrinsic pontine glioma,” Journal of clinical oncology, vol. 28, no. 31, p. 4762, 2010.

[15] E. P. Mitchell, “Oxaliplatin with 5-fu or as a single agent in advanced/metastatic colorectal cancer.” Oncology (Williston Park, NY), vol. 14, no. 12 Suppl 11, pp. 30–32, 2000.

[16] G. L. Lin, K. M. Wilson, M. Ceribelli, B. Z. Stanton, P. J. Woo, S. Kreimer, E. Y. Qin, X. Zhang, J. Lennon, S. Nagaraja et al., “Therapeutic strategies for diffuse midline glioma from high-throughput combination drug screening,” Science translational medicine, vol. 11, no. 519, 2019.

[17] M. Buyl and T. De Bie, “Debayes: a bayesian method for debiasing network embeddings,” arXiv preprint arXiv:2002.11442, 2020.

[18] B. Kang, J. Lijffijt, and T. De Bie, “Conditional network embeddings,” arXiv preprint arXiv:1805.07544, 2018.

[19] P. Sidorov, S. Naulaerts, J. Ariey-Bonnet, E. Pasquier, and P. J. Ballester, “Predicting synergism of cancer drug combinations using nci-almanac data,” Frontiers in Chemistry, vol. 7, 2019.

[20] “Gene ontology resource.” [Online]. Available: http://geneontology.org/

[21] M. Kuhn, I. Letunic, L. J. Jensen, and P. Bork, “The sider database of drugs and side effects,” Nucleic Acids Research, vol. 44, no. D1, 2015.

[22] O. Ursu, J. Holmes, C. G. Bologa, J. J. Yang, S. L. Mathias, V. Stathias, D.-T. Nguyen, S. Schürer, and T. Oprea, “Drugcentral 2018: an update,” Nucleic Acids Research, vol. 47, no. D1, 2018.

[23] T. Sterling and J. J. Irwin, “Zinc 15 – ligand discovery for everyone,” Journal of Chemical Information and Modeling, vol. 55, no. 11, p. 2324–2337, Sep 2015.

[24] X. Yue, Z. Wang, J. Huang, S. Parthasarathy, S. Moosavinasab, Y. Huang, S. M. Lin, W. Zhang, P. Zhang, H. Sun, and et al., “Graph embedding on biomedical networks: methods, applications and evaluations,” Bioinformatics, Apr 2019.

[25] A. Grover and J. Leskovec, “node2vec,” Proceedings of the 22nd ACM SIGKDD International Conference on Knowledge Discovery and Data Mining, 2016.

[26] L. Deng, B. Zou, W. Zhang, and H. Liu, “Drugcombdb: a comprehensive database of drug combinations toward network medicine and combination therapy,” 2018.

[27] M. Adenot and R. Lahana, “Blood-brain barrier permeation models: Discriminating between potential cns and non-cns drugs including p-glycoprotein substrates,” Journal of Chemical Information and Computer Sciences, vol. 44, no. 1, pp. 239–248, 2004, pMID: 14741033. [Online]. Available: https://doi.org/10.1021/ci034205d

[28] Z. Gao, Y. Chen, X. Cai, and R. Xu, “Predict drug permeability to blood–brain-barrier from clinical phenotypes: drug side effects and drug indications,” Bioinformatics, 2016.

[29] A. Ianevski, A. K. Giri, and T. Aittokallio, “Synergyfinder 2.0: visual analytics of multi-drug combination synergies,” Nucleic Acids Research, 2020.

[30] B. Yadav, K. Wennerberg, T. Aittokallio, and J. Tang, “Searching for drug synergy in complex dose–response landscapes using an interaction potency model,” Computational and structural biotechnology journal, vol. 13, pp. 504–513, 2015.

